# Excitatory signal processing in early visual areas based on task-relevance and perceptual salience modulates visual perceptual learning

**DOI:** 10.1101/2025.07.21.665859

**Authors:** Markus Becker, Savanna Babu, Zhiyan Wang, Sebastian M. Frank

## Abstract

Adaptive visual perceptual learning (VPL) should occur primarily for visual signals relevant to an observer’s task but not for task-irrelevant signals. However, it is unclear which neural mechanisms reduce VPL for task-irrelevant signals. Here, we repeatedly exposed participants to a task-irrelevant visual signal (coherent motion in one direction) that was perceptually salient (suprathreshold for coherent motion detection) or weak (near threshold for coherent motion detection). The processing of the task-irrelevant signal in early visual areas during the first and final exposure sessions was measured using functional magnetic resonance spectroscopy (fMRS). The behavioral results showed that discrimination sensitivity for the exposed coherent motion direction increased to a greater extent after near threshold than after suprathreshold exposure. The fMRS results showed that excitatory processing in early visual areas, as reflected in the concentration of glutamate, was lower during suprathreshold than near threshold exposure. The lower the level of excitatory processing in the first exposure session, the more deteriorated the discrimination sensitivity for the coherent motion direction after suprathreshold exposure. When the coherent motion direction was rendered task-relevant, levels of glutamate in early visual areas reversed such that excitatory processing was greater when the coherent motion direction was presented suprathreshold than near threshold. This suggests that the level of excitatory processing in early visual areas changes with the task relevance and perceptual salience of a visual signal, which may modulate the extent of VPL for this signal when it is repeatedly exposed.

**Significance Statement:** Visual perceptual learning (VPL) enhances the efficiency in processing visual signals and is fundamental to the brain’s ability to adapt to changing environmental conditions. However, little is known about neural mechanisms that reduce VPL, although such mechanisms are crucial for adaptive VPL, which occurs primarily for visual signals relevant to an observer’s task but not for task-irrelevant signals. Here we find that early visual areas decrease excitatory processing for a salient task-irrelevant visual signal, which is associated with reduced VPL for this signal. This suggests, first, that a decrease of excitatory activity may be a mechanism that reduces VPL for salient task-irrelevant signals and, second, that this mechanism exerts its effect already at an early stage of cortical visual processing.

## Introduction

A fundamental brain function is the ability to adapt to sensory signals in the environment and prior sensory experiences. An example of this is visual perceptual learning (VPL), which refers to a performance change in a visual task with visual training or repeated visual experience (Dosher & Lu, 2017; Sagi, 2011; Seitz & Dinse, 2007; Watanabe & Sasaki, 2015). Several mechanisms have been identified that facilitate VPL (Censor et al., 2012; Gilbert & Sigman, 2007; Li, 2016; Li et al., 2004; Roelfsema et al., 2010; Sagi, 2011; Sasaki et al., 2010; Seitz & Dinse, 2007; Seitz & Watanabe, 2005; Shibata et al., 2014). However, significantly less is known about mechanisms that reduce VPL. Yet, these mechanisms are crucial for adaptive VPL, which occurs primarily for visual signals relevant to an observer’s task but not for task-irrelevant signals. Task-irrelevant signals are potentially distracting, may interfere with the processing and learning of task-relevant signals, and jeopardize the stability of previously acquired VPL for task-relevant signals (Chang et al., 2014; H. Choi et al., 2009; Egeth & Yantis, 1997; Friedman-Hill et al., 2003; Yan et al., 2014).

Previous studies found that VPL was more pronounced for perceptually weak task-irrelevant signals (Galliussi et al., 2018; Gutnisky et al., 2009; Seitz & Watanabe, 2003; Watanabe et al., 2001) than for perceptually salient task-irrelevant signals (Ahissar & Hochstein, 1993; Chang et al., 2014; Frank, Forster, et al., 2021; Shiu & Pashler, 1992; Tsushima et al., 2008; Vidnyánszky & Sohn, 2005). Based on these results, it was proposed that task-irrelevant signals are detected and suppressed by control mechanisms only if they are sufficiently salient, whereas weak task-irrelevant signals escape this suppression, resulting in VPL for these signals with repeated exposure (Seitz & Watanabe, 2009). However, at what stage in visual processing this suppression of salient task-irrelevant signals occurs, and which neuronal mechanisms are involved has remained uncertain.

Functional magnetic resonance imaging (fMRI) studies found decreased hemodynamic responses in higher visual areas including motion-sensitive area MT+ for salient task-irrelevant coherent motion (Gál et al., 2009; Tsushima et al., 2006), suggesting that the suppression occurs at later stages in visual processing. However, the suppression would be more effective if neuronal responses for task-irrelevant signals were already reduced in early visual areas, including V1, to minimize further processing in other areas. Recent studies suggest that early visual areas may function as ‘adaptive processors’ (Gilbert & Sigman, 2007) whose activities can dynamically adapt to task demands and contexts (Ekstrom et al., 2008; Keller et al., 2017; Li et al., 2004; Ramalingam et al., 2013; Saproo & Serences, 2014; Silvanto et al., 2005; Yang et al., 2024). As such, less pronounced processing of task-irrelevant signals in early visual areas could reduce information flow and further processing of these signals in other visual areas (Kamiyama et al., 2016).

Here, we tested the hypothesis that salient task-irrelevant visual signals are already processed to a lesser extent in early visual areas, reducing VPL for these signals. Participants were repeatedly exposed to a task-irrelevant visual signal (coherent motion in one direction) within an annulus surrounding the screen center. The exposed task-irrelevant signal was either perceptually salient (suprathreshold for coherent motion detection) or weak (near threshold for coherent motion detection) in different groups of participants. The first and final exposure sessions were carried out in the MRI scanner and processing of the task-irrelevant signal was measured in early visual areas using functional magnetic resonance spectroscopy (fMRS). We predicted reduced VPL for the suprathreshold compared with the near threshold task-irrelevant signal and less pronounced processing of this signal in early visual areas during suprathreshold than near-threshold exposure. Moreover, we hypothesized that if early visual areas functioned as adaptive processors, they would reverse the level of processing when the exposed signal was made task-relevant and would exhibit more pronounced processing for a salient than a weak signal.

## Methods

### Participants

A combined total of 82 participants [52 females, 30 males, mean ± standard-error-of-the-mean (SEM) age = 23.70 ± 0.59 years old; 79 right-handed, 3 left-handed] with normal or corrected-to-normal vision was recruited. Participants had normal color-vision as determined by the Ishihara color test. Participants gave informed written consent prior to participation. The study was approved by the internal review board of the University of Regensburg. Thirty participants took part in the behavioral experiment. Thirty new participants took part in the imaging experiment. Another thirty participants took part in the control imaging experiment (fifteen participants for each condition). Of these, twenty-two were new participants. Three participants completed the first condition of the control imaging experiment prior to performing the imaging experiment. Five participants completed the second condition of the control imaging experiment after the imaging experiment.

### Coherent Motion Detection Threshold

The coherent motion detection threshold was measured following previous descriptions (Chang et al., 2014; Frank, Bründl, et al., 2021). On each trial, 70 white dots with a diameter of 0.3° each were presented in a circular field subtending 2.5° to 8° starting from the center of the screen. In the screen center a fixation cross was presented within a gray circle with a diameter of 4°. There were two conditions, coherent motion (signal dots) intermixed with random motion (noise dots) and pure random motion (only noise dots). For coherent motion 3%, 13%, 23%, 33% or 53% of all dots moved together with a speed of 24°/s in one of six directions (10°, 70°, 130°, 190°, 250° or 310°, rotated clockwise from the upward direction). Randomly moving dots were re-displayed at a random location on the screen on every screen flip (limited lifetime = 16.7 ms). Each trial lasted 500 ms. Participants were asked to maintain central fixation. After stimulus offset participants indicated by pressing one of two keys on the keyboard whether they detected coherent motion. No feedback about the correctness of the response was provided. There was a total of 300 trials (150 trials with coherent motion and 150 trials with pure random motion). Each of the five coherence levels was presented 30 times. Trials were presented in random order. For trials with coherent motion, a psychometric function was fitted to the percentage of correct responses averaged across the six different directions. The 80% threshold was used as the individual threshold for coherent motion detection.

### Direction Discrimination Task

On each trial, coherent motion was presented at 0.3 x the individual detection threshold level for 500 ms using the same parameters as in the coherent motion detection threshold measurement. Participants were asked to maintain central fixation and to indicate at the end of the trial in which direction the coherent motion took place. For this purpose, six arrows were displayed around the fixation cross for the six possible coherent motion directions after stimulus offset. Participants selected the coherent motion direction by clicking on the corresponding arrow with the cursor. No feedback about the correctness of the response was provided. There was a total of 120 trials (20 for each coherent motion direction).

### Training Task

Participants performed a rapid-serial-visual-presentation (RSVP) task at the center of the screen with simultaneous exposure to task-irrelevant coherent motion in one direction (Watanabe et al., 2001) (Figure 1). On each 4 s-long trial two images of different animals as targets were presented intermixed with six images of different vegetables, fruits and flowering plants as distractors in the RSVP (following the RSVP design in a previous study; Frank, Bründl, et al., 2021). The two targets were randomly selected from a set of four animal images: a red bird, a yellow fish, a green butterfly, and a blue beetle. The distractors were sourced from three categories, each with eight different images: vegetables (e.g., tomato, corn, broccoli, eggplant), fruits (e.g., apple, banana, kiwi, grape) and flowering plants (e.g., hibiscus, chrysanthemums, clover, forget-me-nots). The distractors were selected to be similar in color to the targets, meaning that there were two red, two yellow, two green and two blue images per category. The images were downloaded from the open-source image database “Pixabay” (https://pixabay.com). The original image background was removed. Images were displayed within a gray circle in the center of the screen with a diameter of 4°. On each trial, target and distractor images were randomly selected from the pool of images and presented in random order. Each image was overlaid with a mask of randomly colored pixels, which replaced 81% of the original image pixels (for illustrative purposes not shown in Figure 1). Each image was presented for 350 ms, preceded and succeeded by on and off ramps with a fixation cross presented within the central gray circle for 75 ms each. At the end of each trial, the images of the four animal targets were displayed side by side on the screen without masks. Participants were asked to report the two animals in the order in which they were presented during the trial by pressing two of four buttons on the button box in the scanner or keys on the keyboard outside the scanner. During behavioral training outside the scanner, feedback about the correctness of participants’ responses was provided by framing the selected animal images in green for a correct response or in red for an incorrect response. Participants were given 2 s to respond. A total of 100 trials was included in each behavioral training session.

**Figure 1.**
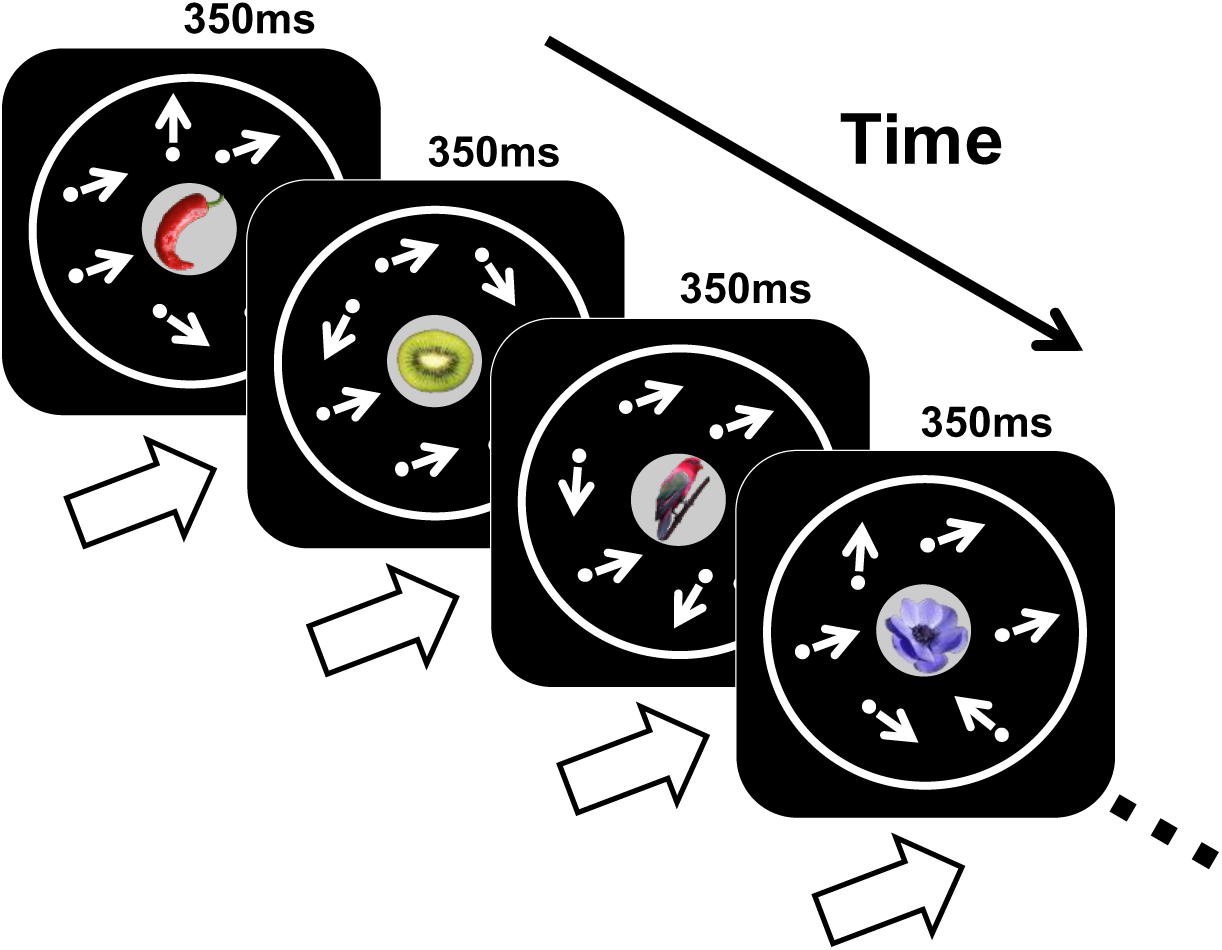
Example trial of the training task. A rapid serial visual presentation (RSVP) stream consisting of images of animals, fruits, vegetables, and flowering plants was presented in the center of the screen. Each image was overlaid with a mask of differently colored pixels (for illustrative purposes not shown). Participants’ task was to detect which animals were presented in which order. Together with the RSVP, moving dots were exposed in the visual periphery. Each dot’s moving direction is illustrated by a small arrow (not shown in the real experiment). A subset of dots moved coherently in the same direction (corresponding to the task-irrelevant learning direction). The arrow below each image shows the task-irrelevant learning direction.

While performing the RSVP task, participants were exposed to task-irrelevant coherent motion in one direction within a circular field subtending 2.5° to 8° starting from the center of the screen (Figure 1). The dot motion parameters were exactly the same as in the coherent motion detection threshold measurement and the direction discrimination task. For each participant one out of six coherent motion directions was randomly selected and consistently exposed together with the RSVP on each trial and each training session (henceforth referred to as the “task-irrelevant learning direction”). There were two exposure groups in the behavioral experiment and in the imaging experiment: the “near threshold exposure group” and the “suprathreshold exposure group”. Depending on the group membership, the salience of the exposed coherent motion direction in the training task was varied. Participants in the near threshold exposure group were exposed to a coherent motion direction near detection threshold for coherent motion (1.0 x individual threshold). Participants in the suprathreshold group were exposed to a coherent motion direction above detection threshold for coherent motion (4.0 x individual threshold) (similar to Chang et al., 2014; Frank, Bründl, et al., 2021). Participants were randomly assigned to one exposure group. They were instructed to maintain central fixation and perform the RSVP. There was no task regarding the peripherally exposed coherent motion.

### Behavioral Experiment

The experiment consisted of a total of eight sessions on separate days (Figure 2A). Participants first completed the coherent motion detection threshold measurement, followed by a Pre-Test in the direction discrimination task (session 1). The mean ± SEM coherent motion detection threshold across participants was 16.8 ± 2.00% in the near threshold exposure group and 17.8 ± 1.71% in the suprathreshold exposure group. Independent sample *t*-tests showed no significant differences in coherent motion detection threshold and Pre-Test discrimination sensitivity of the task-irrelevant learning direction between groups [detection threshold: *t*(28) = -0.36, *p* = 0.72; discrimination sensitivity: *t*(28) = -0.44, *p* = 0.67] (Figure 2B). Participants then completed six sessions in the training task (sessions 2-7). A subset of 9 participants in the near threshold exposure group and a subset of 10 participants in the suprathreshold exposure group performed the training with eye-tracking. For this purpose, the vertical and horizontal position of the right eye was sampled with a frequency of 250 Hz in each trial using a video-based eye-tracking system (Cambridge Research Systems, Kent, UK). Finally, participants completed a Post-Test in the direction discrimination task (session 8). The number of participants for the behavioral experiment was determined based on previous behavioral studies using similar training tasks with comparison between near threshold and suprathreshold exposure groups (Chang et al., 2014; Frank, Bründl, et al., 2021; Tsushima et al., 2008). For each exposure group, seven (Tsushima et al., 2008) and ten participants (Chang et al., 2014; Frank, Bründl, et al., 2021) were recruited in these previous studies. To be sufficiently powered we decided to recruit 15 participants for each exposure group in the behavioral experiment. The results of the behavioral experiment were used to determine the sample size for the imaging experiment. Partial η^2^ for the within-between interaction in a mixed design ANOVA with the within-factor of test (Pre-Test, Post-Test) and the between-factor of exposure group (near threshold, suprathreshold) was calculated (partial η^2^ = 0.20, see *Behavioral Experiment* in *Results* section) and submitted to a power analysis using G*Power (Faul et al., 2007; α = 0.005, 1-β = 0.95; correlation among repeated measures: 0.5). The results showed that the estimated required total sample size was 26 participants. To keep the number of participants similar across the behavioral and imaging experiment we decided to recruit a total of 30 new participants (15 for each exposure group) for the imaging experiment.

**Figure 2.**
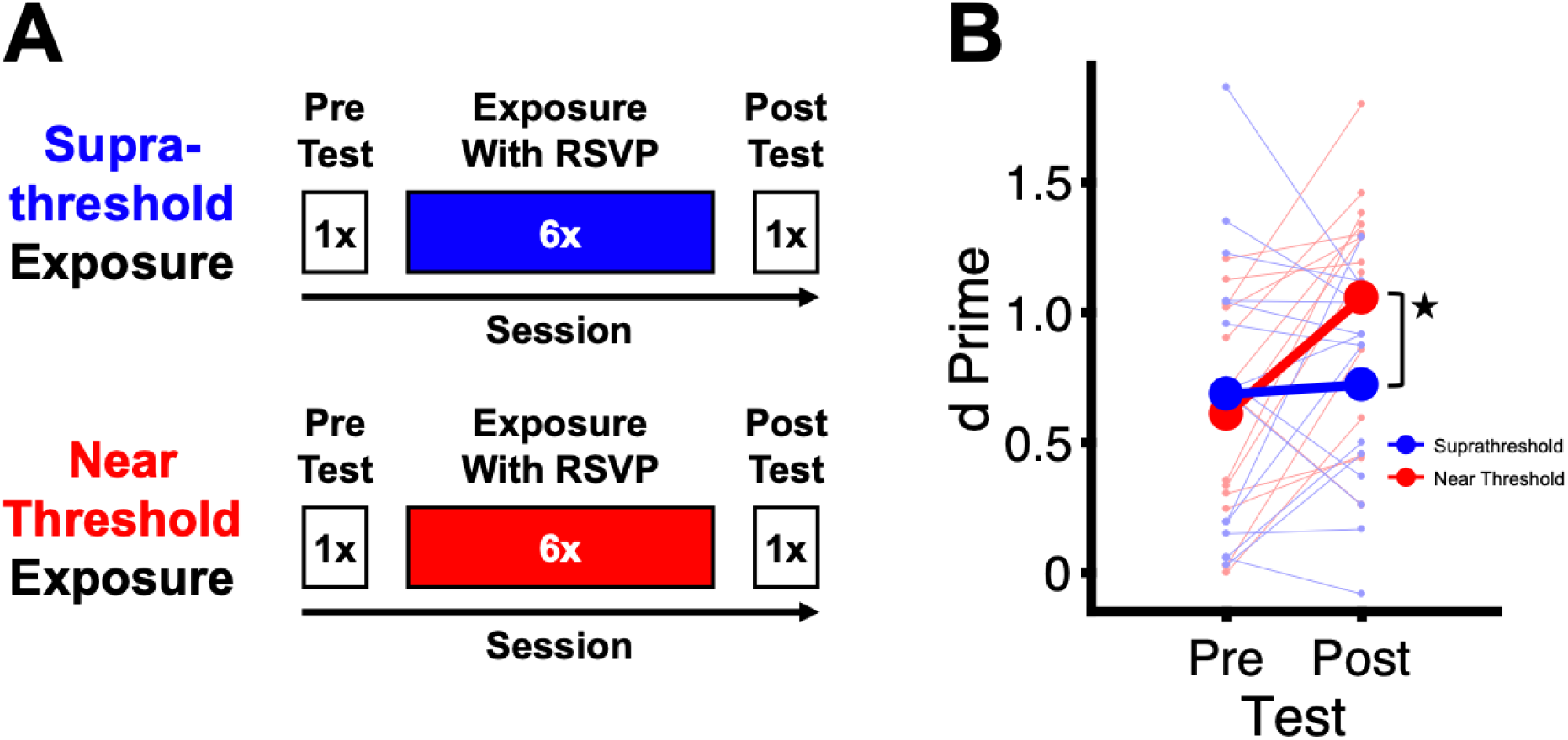
Design and results of the behavioral experiment. (**A**) Experimental design. Participants in the near threshold group were exposed to the task-irrelevant learning direction near detection threshold for coherent motion. Participants in the suprathreshold group were exposed to the task-irrelevant learning direction above detection threshold for coherent motion. (**B**) Discrimination sensitivity (d’) for the exposed task-irrelevant learning direction prior to the first exposure session (Pre-Test) and after the final exposure session (Post-Test). Thick lines show mean results across participants in each exposure group. Thin lines show results for each participant. The asterisk shows a significant interaction between test and exposure group. * *p* < 0.05.

### Imaging Experiment

The imaging experiment consisted of a total of ten sessions on separate days (Figure 3A). In the first session participants performed the coherent motion detection threshold measurement followed by a Pre-Test in the direction discrimination task outside the scanner. The mean ± SEM coherent motion detection threshold across participants was 15.0 ± 0.94% in the near threshold exposure group and 13.9 ± 1.19% in the suprathreshold exposure group. Independent sample *t*-tests showed no significant differences in coherent motion detection threshold and Pre-Test discrimination sensitivity of the task-irrelevant learning direction between groups [detection threshold: *t*(28) = 0.69, *p* = 0.50; discrimination sensitivity: *t*(28) = -0.14, *p* = 0.89] (Figure 3B). In the second session participants performed the training task in the MRI scanner (Pre-MRI). Then, participants completed six sessions in the training task outside the scanner exactly as in the behavioral experiment (sessions 3 to 8). In the ninth session participants performed the training task again in the MRI scanner (Post-MRI). Two participants in the near threshold exposure group were not available for the Post-MRI scan. Finally, in the tenth session, participants completed a Post-Test in the direction discrimination task exactly as the Pre-Test outside the scanner. The two MRI sessions took place inside a Prisma 3 Tesla MRI scanner (Siemens, Erlangen, Germany) using a head/neck coil with 64 channels. In each MRI session single-voxel proton (^1^H) fMRS measurements were conducted for a volume-of-interest (VOI) in early visual areas while participants performed the training task and were exposed to the task-irrelevant learning direction. For fMRS, a MEGA-PRESS sequence (Mescher et al., 1996, 1998) with alternating Edit On and Edit Off scans [time-to-repeat (TR) = 1.5 s; time-to-echo (TE) = 68 ms; flip angle (FA) = 90°; number of averages = 128] was used. A frequency selective, single band Gauss pulse was utilized to saturate the *β*-CH_2_ signal at 1.94 ppm and to refocus the J evolution of the triplet γ-CH_2_ resonance of γ-aminobutyric acid (GABA) at 3 ppm (‘Edit On’). The very same Gauss pulse was used to irradiate the opposite part of the spectrum at 7.46 ppm (‘Edit Off’). The ‘Edit Off’ spectrum was subtracted from the ‘Edit On’ spectrum to produce a difference spectrum. WET water suppression was used (Ogg et al., 1994). The VOI had a size of 2 x 2 x 2 cm. It was placed perpendicular to the calcarine sulcus and centered between the left and right occipital lobes, maintaining some distance from the occipital poles (see Figure 4A). Non-neural tissue containing lipids and proximity to the dura were avoided during VOI placement. For the purpose of VOI placement three short high-resolution anatomical scans for the coronal plane [TR = 0.25 s, TE = 2.46 ms, FA = 70°, in-plane acquisition matrix (AM) = 288 x 288, 35 slices, voxel size = 0.8 x 0.8 x 4.0 mm, inter-slice gap = 1.20 mm], sagittal plane (TR = 0.19 s, TE = 2.46 ms, FA = 70°, AM = 288 x 288, 25 slices, voxel size = 0.8 x 0.8 x 4.0 mm, inter-slice gap = 1.20 mm) and transverse plane (TR = 0.19 s, TE = 2.46 ms, FA = 70°, AM = 288 x 288, 27 slices, voxel size = 0.8 x 0.8 x 4.0 mm, inter-slice gap = 1.20 mm) were collected prior to fMRS. Furthermore, a water reference scan for the same VOI was collected after fMRS using a PRESS sequence without water suppression (TR = 3 s; TE = 30 ms; FA = 90°; number of averages = 8). A checklist containing all fMRS acquisition parameters, analyses and quality checks, as proposed by Choi et al. (2021), is provided in Supplementary Table 1.

**Figure 3.**
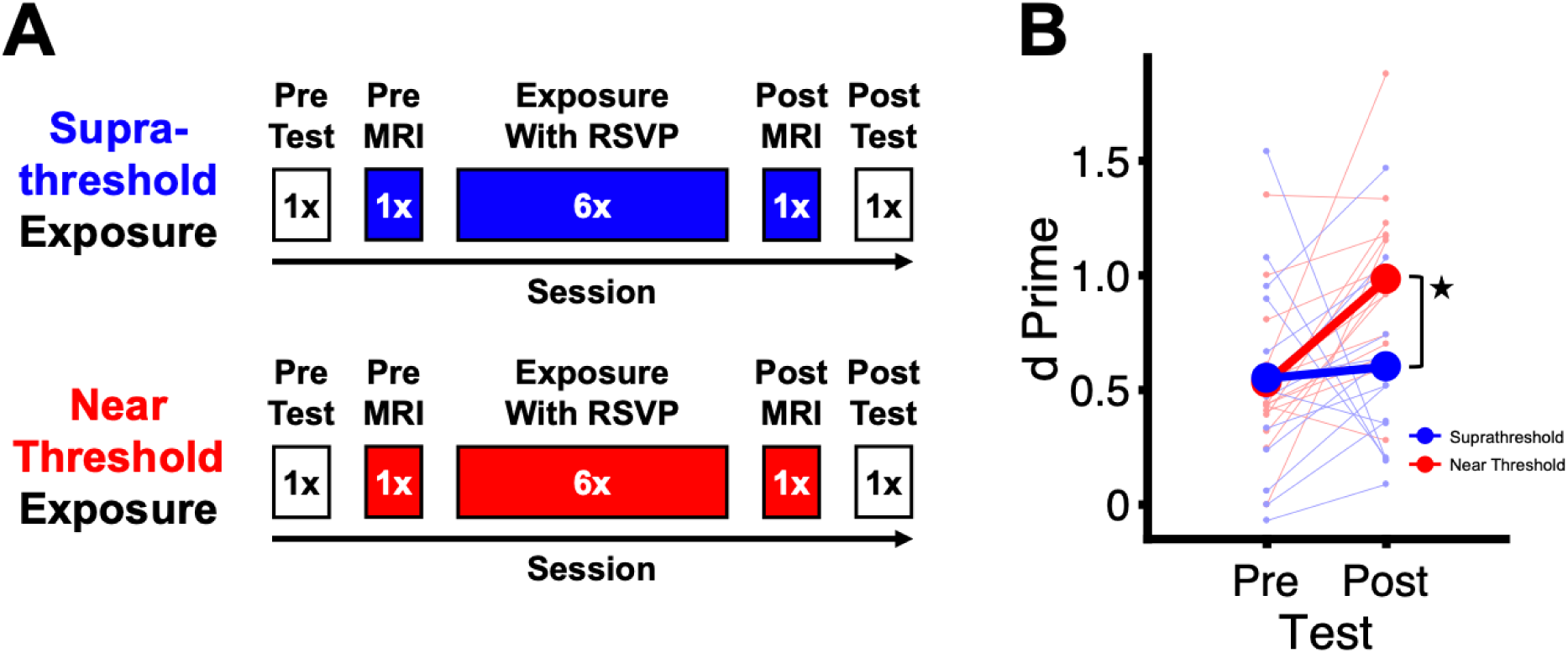
Design and behavioral results of the imaging experiment. (**A**) Experimental design. The first and final exposure sessions (Pre-MRI and Post-MRI) in each group were conducted in the scanner. Otherwise same as Figure 2A. (**B**) Behavioral results. Otherwise same as Figure 2B. * *p* < 0.05.

**Figure 4.**
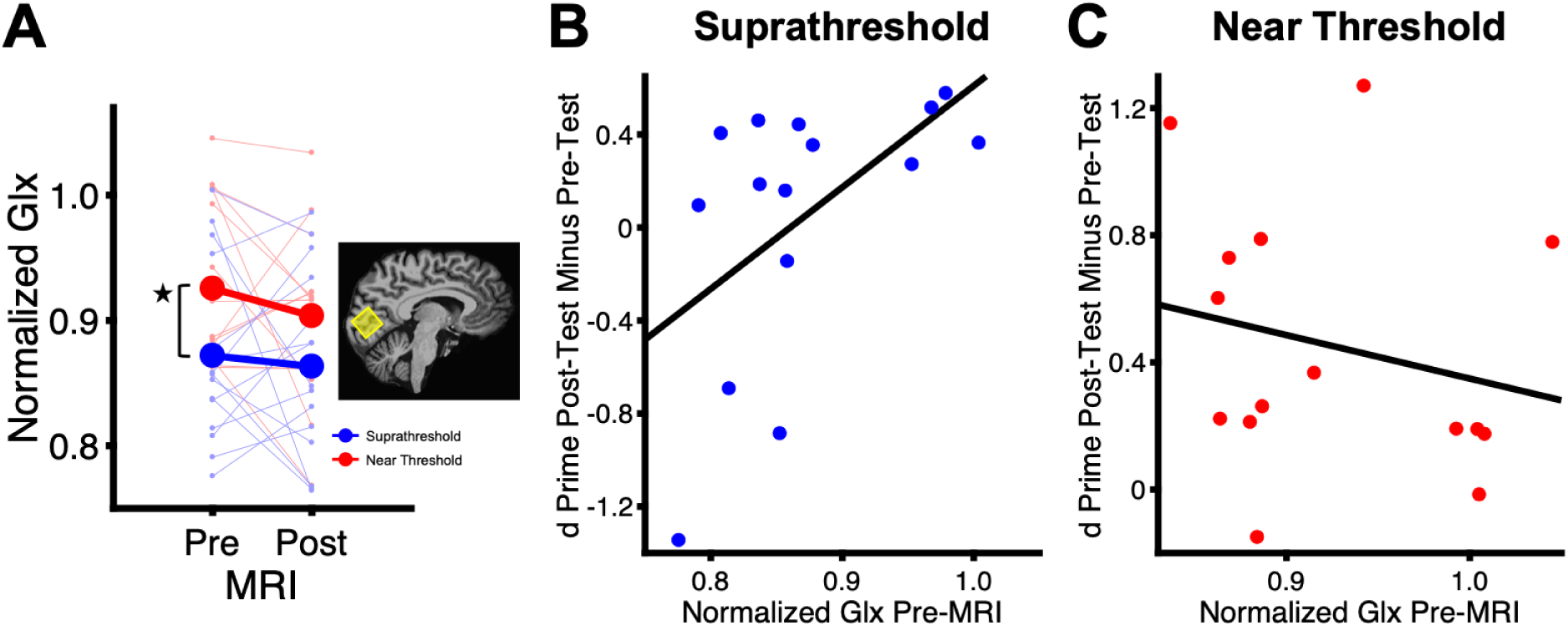
Excitatory processing levels in early visual areas in the imaging experiment. (**A**) Glx concentration normalized to a control metabolite. Greater values on the y-axis correspond to greater Glx concentration. The inset shows the location of the fMRS volume-of-interest in a representative participant. Thick lines show mean results across participants in each exposure group. Thin lines show results for each participant. The asterisk shows a significant main effect of exposure group. * *p* < 0.05. (**B**) Correlation between Glx concentration in Pre-MRI and sensitivity change for the task-irrelevant learning direction from Pre-Test to Post-Test (see Figure 3B) in the suprathreshold exposure group. Each dot shows the result from a different participant in the suprathreshold exposure group. (**C**) Correlation results in the near threshold exposure group. Otherwise same as (B).

One fMRS run was conducted in each MRI session during which participants completed 70 trials in the training task with exposure to the task-irrelevant learning direction. Trials were presented continuously without intermission. No feedback about response accuracy in the RSVP was provided in the scanner. The response options selected by participants at trial end were framed in neutral blue. The mean ± SEM percentage overlap in VOI location between Pre-MRI and Post-MRI across participants was 74.8 ± 3.57% (near threshold exposure group) and 78.3 ± 2.38% (suprathreshold exposure group). Automatic shimming (gradient field adjustments to increase the homogeneity of the magnet field B_0_) was conducted prior to fMRS. For each participant the shim values were kept below a full-width at half-maximum (FWHM) of 20 Hz. The mean ± SEM shim values across participants were as follows: 13.9 ± 0.25 Hz in Pre-MRI and 13.9 ± 0.20 Hz in Post-MRI (near threshold exposure group); 13.3 ± 0.21 Hz in Pre-MRI and 13.5 ± 0.17 Hz in Post-MRI (suprathreshold exposure group). In Pre-MRI, a high-resolution anatomical scan of the brain was collected using a magnetization prepared rapid gradient echo sequence (TR = 2.3 s, TE = 2.32 ms, FA = 8°, AM = 256 x 256, 192 sagittal slices, voxel size = 0.9 x 0.9 x 0.9 mm, interslice gap = 0.45 mm).

### Control Imaging Experiment

The aim of the control imaging experiment was to examine whether differences in glutamatergic processing in early visual areas between near threshold and suprathreshold coherent motion is modulated by task-relevance. For this purpose, two conditions were carried out in different groups of participants. In the first condition the coherent motion was task-irrelevant as in the imaging experiment. Participants in this condition first completed a coherent motion detection threshold measurement outside the scanner. The mean ± SEM coherent motion detection threshold across participants was 14.0 ± 0.93%. Then, they performed two runs in the training task (one run with exposure to task-irrelevant near threshold coherent motion and another run with exposure to task-irrelevant suprathreshold coherent motion) while fMRS was measured exactly as in the imaging experiment. The run order was counterbalanced across participants. One coherent motion direction was randomly selected for each participant and presented at near threshold and at suprathreshold levels. The selected direction was counterbalanced across participants. After each fMRS run a water reference scan was collected exactly as in the imaging experiment. The mean ± SEM RSVP response accuracy across participants was 58.3 ± 5.04% correct in near threshold exposure and 56.8 ± 6.21% correct in suprathreshold exposure. The mean ± SEM shim values across participants were 13.6 ± 0.29 Hz for near threshold exposure and 13.5 ± 0.27 Hz for suprathreshold exposure.

In the second condition the peripheral coherent motion was rendered task-relevant. Participants first completed a coherent motion detection threshold measurement and a practice session in a modified version of the training task outside the scanner. The mean ± SEM coherent motion detection threshold across participants was 16.4 ± 0.80%. The training task was modified in the following way: first, the mask overlaid on the images in the central RSVP was weakened by replacing only 50% of the original image pixels with randomly colored pixels. With this weaker mask, untrained participants were able to perform the RSVP task with high response accuracy (∼ 90% correct response accuracy in the RSVP without any training). Second, the peripheral stimulus was rendered task-relevant by asking participants to perform a direction change detection task on the coherent motion. For this purpose, the coherent motion direction changed briefly for 350 ms from the displayed direction. This change in coherent motion direction occurred unpredictably one to three times or not at all during a trial. There was a minimum interval of 300 ms between two successive changes in coherent motion direction and no change occurred in the first and final 150 ms of a trial. Third, participants performed two tasks: at trial end, they first reported which two target animals they had detected in the RSVP in the order of presentation (2 s response time) and then how many changes in the coherent motion direction they had detected (either one, two, three or none; again 2 s response time). Similar to the response options for the RSVP, the four response options for the coherent motion direction change detection task were displayed side by side and participants responded by pressing one of four keys on the keyboard or one of four buttons on the button box in the scanner. No feedback about response accuracy was provided. The selected response options were framed in neutral blue for each task. Participants completed two runs in which the peripheral coherent motion was presented near threshold and suprathreshold for coherent motion detection, respectively. Each run included a total of 50 trials. One coherent motion direction was randomly selected for each participant (presented at near threshold and at suprathreshold levels). The selected direction was counterbalanced across participants. During the practice session, the change in coherent motion direction required to achieve 50% or greater response accuracy in the coherent motion direction change detection task while simultaneously performing the RSVP was determined for each participant. The mean ± SEM change across participants was 39.5 ± 2.55° (near threshold coherent motion) and 12.9 ± 2.76° (suprathreshold coherent motion). The mean ± SEM response accuracy across participants was 95.2% ± 1.09% in the RSVP task and 52.16% ± 1.94% in the peripheral task. After the practice session in the modified version of the training task outside the scanner, participants performed the same task in the scanner while fMRS was measured exactly as in the imaging experiment. After each fMRS run a water reference scan was collected. The mean ± SEM response accuracy across participants was 97.0% ± 0.52% in the RSVP task and 52.9% ± 2.55% in the peripheral task. The mean ± SEM shim values across participants were as follows: 13.9 ± 0.28 Hz (near threshold coherent motion) and 14.0 ± 0.30 Hz (suprathreshold coherent motion).

### Stimulus Presentation and Response Collection

Stimuli were generated using Psychtoolbox (Version 3.0.19; Brainard, 1997; Pelli, 1997) running in Matlab (The Mathworks, Natick, MA, USA). Outside the scanner, stimuli were presented on a gamma-corrected LCD screen with a refresh rate of 60 Hz and a viewing distance of 60 cm in a behavioral testing room with lights turned off. Participants used a chin rest and responded by pressing a key on the keyboard with their right hand. In the scanner, stimuli were presented on a translucent screen located at the back of the MRI bore using a projector with gamma correction and a refresh rate of 60 Hz. Participants viewed the screen with a headcoil-mounted mirror (viewing distance = 97 cm). Participants responded by button press on the MRI-safe button box placed in their right hand.

### Behavioral Analysis

Discrimination sensitivity of the task-irrelevant learning direction was calculated as d’. The RSVP performance was calculated as response accuracy (number of RSVP trials with correct response relative to the total number of RSVP trials per behavioral session or fMRS run in percent). An RSVP trial was scored as correct when the following two conditions were met: first, the participant correctly selected the two target images, and second, the participant selected the two target images in the correct order of presentation. If the two conditions were not met or if the participant did not respond, the trial was scored as incorrect.

### Eye-Tracking Analysis

In the behavioral experiment, the percentage of time with fixation on the RSVP was calculated for each trial in each exposure session and for each participant. Then, for each participant, a linear fit on percentage of time with fixation on the RSVP across exposure sessions was calculated and the slope of this fit was compared with zero (corresponding to no change across exposure sessions) across participants.

### Imaging Analysis

The fMRS data were analyzed using the LC Model (Provencher, 1993, 2001) in the chemical shift range between 1.7 and 4.0 ppm for the Edit Off spectrum and between 1.95 and 4.2 ppm for the difference spectrum (following the approach used by Maddock et al., 2018). For spectra see Supplementary Figures 5 and 6. Water scaling and eddy current correction were performed. The metabolite intensities were fitted to a linear combination of spectra of individual metabolites derived from imported metabolite basis sets (Provencher, 1993, 2001). The concentration of the composite measure of glutamate and its precursor glutamine (together referred to as Glx) was calculated from the Edit Off-spectrum. The concentration of GABA was calculated from the difference spectrum (Maddock et al., 2018). Glx and GABA concentrations were normalized to the composite measure of N-Acetylaspartate and N-Acetylaspartylglutamate (henceforth together referred to as NAA). N-Acetylaspartate is the hydrolysis product of N-Acetylaspartylglutamate and a marker of neuronal density and mitochondrial function (Schuff et al., 2006). NAA is commonly used as a control metabolite to normalize the concentrations of Glx and GABA (Duncan et al., 2014; Frank et al., 2023; Mullins et al., 2014; Stagg, 2014). For the normalization of Glx, the NAA concentration was calculated from the Edit Off spectrum. For the normalization of GABA, the NAA concentration was calculated from the difference spectrum. Only spectral results with a signal-to-noise (SN) ratio greater than 5% as estimated by the LC model were included (see Wilson et al., 2019). No spectrum in any experiment had to be excluded. The mean SNs (± SEM) across participants and experiments were as follows: 33.1 ± 0.28% (Edit Off spectrum) and 28.8 ± 0.93% (difference spectrum).

The Cramer-Rao Lower Bounds (CRLB) indicate the reliability of quantification of Glx, GABA and NAA. A criterion of 25% was used to reject low-quality results (Frank et al., 2022). No scans for Glx or NAA had to be excluded in any experiment. GABA results from one participant in the suprathreshold exposure group in Pre-MRI in the imaging experiment had to be excluded. The mean CRLB ± SEM across participants and experiments were as follows: 7.78 ± 0.13% for Glx (Edit Off spectrum), 2.15 ± 0.05% for NAA (Edit Off spectrum), 17.3 ± 0.46% for GABA (difference spectrum), and 3.15 ± 0.06% for NAA (difference spectrum).

Prior to normalization the concentrations of Glx, GABA, and NAA were further corrected for each participant’s volume fractions inside the fMRS VOI. For this purpose, volume fractions within the fMRS VOI corresponding to gray matter, white matter and other types of volume including cerebrospinal fluid were calculated using the Freesurfer segmentation of each participant’s reconstructed high-resolution anatomical scan of the brain (Freesurfer Version 7.3.2; Martinos Center for Biomedical Imaging, Charlestown, MA, USA; Dale et al., 1999; Fischl et al., 1999). The mean ± SEM volume fractions across participants were 53.1 ± 0.37% (gray matter), 37.1 ± 0.35% (white matter) and 9.79 ± 0.19% (other). The same correction approach as in Kolasinski et al. (2017) was used. Glx and GABA were corrected for the proportion of gray matter within the VOI by dividing by [gray matter / (gray matter + white matter + other)]. NAA was corrected for the proportion of total brain volume within the VOI by dividing by [(gray matter + white matter) / (gray matter + white matter + other)]. Then, for each participant and fMRS scan the corrected concentrations of Glx and GABA were normalized to the corrected concentration of NAA. Post-hoc analyses showed that within the gray matter of the fMRS VOI 60.3 ± 1.22% corresponded to V1, 28.6 ± 0.87% corresponded to V2, and 3.78 ± 0.38% corresponded to V3 across participants and experiments. Gray matter corresponding to other cortical areas was negligible. Visual areas were defined by remapping the cortical segmentation proposed by Glasser et al. (2016) from the Freesurfer template brain to each participant’s reconstructed brain. The fMRS VOI had the following mean ± SEM MNI coordinates across participants and experiments: X = 1.03 ± 0.21, Y = -79.7 ± 0.43, Z = 7.93 ± 0.73.

### Statistics

Behavioral and imaging results were analyzed using parametric statistics (ANOVA and *t*-test) with a two-tailed alpha-level of 0.05. Partial η^2^, Cohen’s *d*, and *r* were calculated as measures of effect size for ANOVA, *t*-test, and Pearson correlation, respectively. Cohen’s *d* for repeated and non-repeated measures was calculated using the formulas implemented in G*Power (Faul et al., 2007).

## Results

### Behavioral Experiment

Figure 2B shows discrimination sensitivity for the task-irrelevant learning direction in Pre-Test and Post-Test. A 2 x 2 mixed design ANOVA with the within-factor of test (Pre-Test, Post-Test) and the between-factor of exposure group (near threshold, suprathreshold) showed a significant interaction between test and exposure group [*F*(1,28) = 6.78, *p* = 0.01, partial η^2^ = 0.20], suggesting that discrimination sensitivity improved greater from Pre-Test to Post-Test in the near threshold than in the suprathreshold exposure group (Figure 2B). There was a significant main effect of test [*F*(1,28) = 9.31, *p* = 0.005, partial η^2^ = 0.25], indicating that discrimination sensitivity improved from Pre-Test to Post-Test irrespective of exposure group (Figure 2B). There was no significant main effect of exposure group [*F*(1,28) = 0.77, *p* = 0.39]. Post-hoc analyses showed that discrimination sensitivity was not significantly different between exposure groups in Pre-Test [independent sample *t*-test; *t*(28) = -0.44, *p* = 0.67] but significantly greater after near threshold exposure than after suprathreshold exposure in Post-Test [*t*(28) = 2.15, *p* = 0.04, *d* = 0.78].

Participants’ performance on the RSVP task improved from the first to the final exposure session (Supplementary Figure 1). A 2 x 2 mixed design ANOVA with the within-factor of session (first exposure session, final exposure session) and the between-factor of exposure group (near threshold, suprathreshold) showed a significant main effect of session [*F*(1,28) = 50.6, *p* < 0.001, partial η^2^ = 0.64], suggesting that performance in the RSVP task improved from the first to the final exposure session. There was no significant main effect of exposure group [*F*(1,28) = 0.73, *p* = 0.40] and no significant interaction between session and exposure group [*F*(1,28) = 0.01, *p* = 0.91]. The RSVP learning speed, i.e., the slope of improvement, quantified by a logarithmic fit to RSVP performance across exposure sessions for each participant, was not significantly different between the near threshold and suprathreshold exposure groups [independent sample *t*-test: *t*(28) = 0.14, *p* = 0.89]. Slopes of a linear fit to percentage fixation on the RSVP across exposure sessions for each participant were not significantly different form zero [one-sample *t*-test; near threshold group: *t*(8) = 0.43, *p* = 0.68; suprathreshold group: *t*(9) = 0.31, *p* = 0.76] and there was no significant difference in slopes between the near threshold and suprathreshold exposure groups [independent-sample *t*-test; *t*(17) = -0.23, *p* = 0.82] (Supplementary Figure 2).

### Imaging Experiment

A 2 x 2 mixed design ANOVA with the within-factor of test (Pre-Test, Post-Test) and the between-factor of exposure group (near threshold, suprathreshold) showed a significant interaction between test and exposure group [*F*(1,28) = 4.78, *p* = 0.04, partial η^2^ = 0.15], suggesting that discrimination sensitivity improved greater from Pre-Test to Post-Test in the near threshold than in the suprathreshold exposure group (Figure 3B), similar to the results of the behavioral experiment (see Figure 2B). There was a significant main effect of test [*F*(1,28) = 7.48, *p* = 0.01, partial η^2^ = 0.21], indicating that discrimination sensitivity improved from Pre-Test to Post-Test irrespective of exposure group (Figure 3B). There was no significant main effect of exposure group [*F*(1,28) = 2.90, *p* = 0.10]. Post-hoc analyses showed that discrimination sensitivity was not significantly different between exposure groups in Pre-Test [independent sample *t*-test; *t*(28) = -0.14, *p* = 0.89] but significantly greater after near threshold exposure than after suprathreshold exposure in Post-Test [*t*(28) = 2.70, *p* = 0.01, *d* = 0.99].

Participants’ performance on the RSVP task improved from the first to the final exposure session (Supplementary Figure 3). A 2 x 2 mixed design ANOVA with the within-factor of MRI (Pre-MRI, Post-MRI) and the between-factor of exposure group (near threshold, suprathreshold) showed a significant main effect of MRI [*F*(1,26) = 40.0, *p* < 0.001, partial η^2^ = 0.61], suggesting that performance in the RSVP task improved from Pre-MRI to Post-MRI (Supplementary Figure 3). There was no significant main effect of exposure group [*F*(1,26) = 1.00, *p* = 0.33] and no significant interaction between MRI and exposure group [*F*(1,26) = 0.04, *p* = 0.84]. The RSVP learning speed was not significantly different between the near threshold and suprathreshold exposure groups [independent sample *t*-test: *t*(28) = 0.03, *p* = 0.98].

Figure 4 shows excitatory processing levels measured with fMRS. A 2 x 2 mixed design ANOVA with the within-factor of MRI (Pre-MRI, Post-MRI) and the between-factor of exposure group (near threshold, suprathreshold) on normalized Glx concentration during exposure to the task-irrelevant learning direction in early visual areas showed a significant main effect of exposure group [*F*(1,26) = 5.51, *p* = 0.03, partial η^2^ = 0.18], suggesting that the concentration of Glx was greater in the near threshold than in the suprathreshold exposure group across MRI sessions (Figure 4A). There was no significant main effect of MRI [*F*(1,26) = 1.68, *p* = 0.21] and no significant interaction between exposure group and MRI [*F*(1,26) = 0.55, *p* = 0.46]. Post-hoc analyses showed that the concentration of Glx was significantly greater in the near threshold exposure group than in the suprathreshold exposure group in Pre-MRI [independent sample *t*-test; *t*(28) = 2.11, *p* = 0.04, *d* = 0.77]. There was no significant difference between groups in Post-MRI [*t*(26) = 1.45, *p* = 0.16]. The concentration of Glx in Pre-MRI predicted the sensitivity change for the task-irrelevant learning direction after suprathreshold exposure: participants who tended to have lower Glx concentration in Pre-MRI tended to have decreased sensitivity for the task-irrelevant learning direction after suprathreshold exposure (Pearson correlation; *r* = 0.54, *p* = 0.04) (Figure 4B). No significant correlation was found between Glx concentration in Pre-MRI and sensitivity change for the task-irrelevant learning direction after near threshold exposure (*r* = -0.22, *p* = 0.42) (Figure 4C). There were no significant correlations between Glx concentration in Pre-MRI and subsequent RSVP learning speed (suprathreshold exposure group: *r* = -0.14, *p* = 0.62; near threshold exposure group: *r* = -0.16, *p* = 0.57).

A 2 x 2 mixed design ANOVA with the within-factor of MRI (Pre-MRI, Post-MRI) and the between-factor of exposure group (near threshold, suprathreshold) on GABA concentration showed no significant main effects of exposure group [*F*(1,25) = 0.14, *p* = 0.72] and MRI [*F*(1,25) = 0.16, *p* = 0.69] and no significant interaction between exposure group and MRI [*F*(1,25) = 0.17, *p* = 0.68] (Supplementary Figure 4A). No significant correlations were found between GABA concentration in Pre-MRI and sensitivity change for the task-irrelevant learning direction after near threshold exposure (*r* = -0.25, *p* = 0.37) and after suprathreshold exposure (*r* = 0.07, *p* = 0.80) (Supplementary Figure 4B,C).

### Control Imaging Experiment

Figure 5 shows excitatory processing levels in the control imaging experiment when the coherent motion direction was task-irrelevant and task-relevant in different conditions. A 2 x 2 mixed design ANOVA with the within-factor of threshold level (near threshold, suprathreshold) and task-relevance (task-irrelevant, task-relevant) on the concentration of Glx showed a significant interaction between threshold level and task-relevance [*F*(1,28) = 10.8, *p* = 0.003, partial η^2^ = 0.28], suggesting that the concentration of Glx for the near threshold and suprathreshold coherent motion direction was modulated by task-relevance (Figure 5). There were no significant main effects of threshold level [*F*(1,28) = 0.0002, *p* = 0.99] and task-relevance [*F*(1,28) = 2.83, *p* = 0.10]. Post-hoc analyses showed that the concentration of Glx was significantly greater for the near threshold than for the suprathreshold task-irrelevant coherent motion direction [paired sample *t*-test; *t*(14) = 2.52, *p* = 0.02, *d* = 0.65], similar to the results of the imaging experiment (Figure 4A). In the task-relevant condition, the concentration of Glx was significantly greater for the suprathreshold than for the near threshold task-relevant coherent motion direction [paired-sample *t*-test: *t*(14) = 2.16, *p* = 0.049, *d* = 0.56].

**Figure 5.**
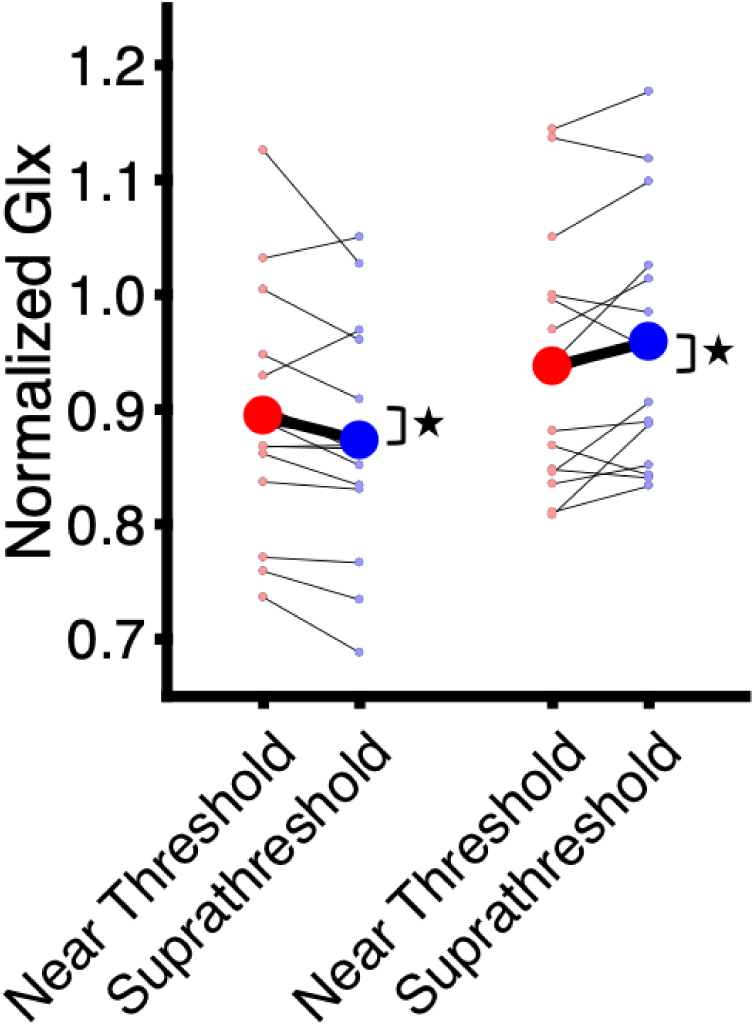
Excitatory processing levels corresponding to Glx concentration normalized to a control metabolite in early visual areas in the control imaging experiment. Large dots show mean results across participants. Small dots show results for each participant. The asterisks show significant differences in Glx concentration between a task-irrelevant (left panel) and task-relevant (right panel) near threshold and suprathreshold coherent motion direction. * *p* < 0.05.

## Discussion

In this study, the role of early visual areas in VPL of task-irrelevant visual signals was examined. Over the course of multiple sessions, participants were repeatedly exposed to a task-irrelevant motion direction either suprathreshold or near threshold for coherent motion detection. VPL corresponded to an increase in discrimination sensitivity for the exposed coherent motion direction after the end of the exposure. Processing of the task-irrelevant coherent motion direction was measured in early visual areas during the first and final exposure sessions using fMRS. The results showed, first, that VPL was significantly more pronounced after near threshold than after suprathreshold exposure, second, that excitatory processing, as reflected in the concentration of glutamate, in early visual areas was significantly lower during suprathreshold than near threshold exposure, and third, that lower levels of excitatory processing predicted greater deterioration of discrimination sensitivity for the coherent motion direction after suprathreshold exposure. The results of a control experiment showed that levels of excitatory processing in early visual areas for near threshold and suprathreshold coherent motion reversed when the motion direction was rendered task-relevant.

The fMRS results indicate that visual signals are processed differently in early visual areas depending on their task relevance and perceptual salience. This difference in processing was reflected in the concentration of glutamate, a chief excitatory neurotransmitter (Petroff, 2002; Reznikov et al., 2011; Sah et al., 2008; Watkins & Jane, 2006). Previous results in animal-models and computational studies found that glutamate plays a crucial role in regulating information flow in the visual processing pathway by modulating the balance of feedforward and feedback communication between the thalamus and cortical areas (Kamiyama et al., 2016; Self et al., 2012). Studies in humans found that levels of glutamate in early visual areas increased with perceptual salience (Ip et al., 2017, 2019; Kurcyus et al., 2018), suggesting that excitatory processing is crucial for signal-driven feedforward visual processing. The results of this study suggest that glutamate levels in early visual areas reflect not only excitatory processing driven by perceptual salience but also the behavioral context of visual signals. For task-irrelevant visual signals, less pronounced excitatory processing in early visual areas could reduce feedforward information flow to other visual areas (Schallmo et al., 2018). Early visual areas process primitive local motion signals, which are integrated over space and time at a later stage in the visual processing pathway in areas such as MT+ (Braddick et al., 2001; Koyama et al., 2005; Movshon & Newsome, 1996; Salzman et al., 1992; Simoncelli & Heeger, 1998; Vaina et al., 2003; Wang et al., 2021; Williams & Sekuler, 1984; Zeki, 1974). Less pronounced excitatory processing in early visual areas, as found in this study, could therefore reflect reduced input of local motion information to higher visual areas (Chen et al., 2015, 2016; Shibata et al., 2016) and also parietal areas involved in decision making about coherent motion directions (Law & Gold, 2008). The amount of further processing of the task-irrelevant visual signal would therefore be reduced and VPL for this signal would be less pronounced.

There are two potential hypotheses that may explain the modulation of glutamatergic processing based on task relevance in early visual areas. One possibility is that the modulation occurs in a top-down fashion through long-range feedback connections from higher areas (Astorga et al., 2022; Gilbert & Li, 2013; Gilbert & Wiesel, 1990; Gutnisky et al., 2009; Ito & Gilbert, 1999; Li et al., 2004; Saproo & Serences, 2014; Tsushima & Watanabe, 2009). In support of this, a previous fMRI study found that participants showed higher lateral prefrontal and lower MT+ blood oxygenation level-dependent responses to salient compared with weak task-irrelevant coherent motion, suggesting that top-down inhibitory control mechanisms influence lower-level visual processing (Tsushima et al., 2006). On the other hand, recent evidence suggests that bottom-up processes could also modulate visual information processing in sensory cortical areas. Arousal states provide strong behavioral context and can directly influence the processing of visual input in the thalamus (McGinley, 2020). Higher levels of arousal could therefore increase processing in early visual areas through increased subcortical input (Liang et al., 2020; Schröder et al., 2020; Vinck et al., 2015). Computational models showed that the basal ganglia and subcortical structures in the midbrain can track the task-relevance of sensory signals and gate information flow from early sensory to other areas by modulating excitatory neural connections (O’Reilly & Frank, 2006). Therefore, it is also possible that glutamatergic processing in early visual areas is modulated through changing bottom-up subcortical input. Further experiments are needed to clarify the contribution of top-down and bottom-up processes to the modulation of excitatory processing in early visual areas depending on task relevance.

Our results showed that participants in the suprathreshold exposure condition either exhibited little improvement in discrimination sensitivity for the exposed coherent motion direction, suggesting reduced VPL, or even tended to deteriorate their discrimination sensitivity for this direction. Previous studies found that repeated exposure to salient task-irrelevant visual signals can lead to deteriorated perception of the exposed signal when it is subsequently tested under task-relevant conditions, potentially reflecting that the processing of the signal was repeatedly suppressed when it was exposed as task-irrelevant (Frank, Bründl, et al., 2021; Vidnyánszky & Sohn, 2005). Suppressing the task-irrelevant visual signal could have the advantage that there is less distraction from the relevant task (e.g., Tsushima et al., 2006), but at the expense of decreased perceptual sensitivity to the suppressed signal if it later has to be processed as task-relevant.

We used fMRS to measure whether signal processing in early visual areas is modulated by task-relevance and perceptual salience. Our fMRS results showing lower levels of glutamatergic processing in association with less pronounced VPL for a salient task-irrelevant visual signal agree with previous studies reporting decreased glutamatergic processing during task-induced or stimulation-induced suppression (Apšvalka et al., 2015; Frank, Forster, et al., 2021; Lally et al., 2014; Stagg et al., 2009). We found no significant modulations of GABA levels with task-relevance and perceptual salience and no significant associations between GABA and VPL. We could speculate, as discussed above, that the decrease of glutamatergic processing for the suprathreshold task-irrelevant coherent motion direction may result from reduced subcortical modulatory input to early visual areas without involving cortical changes in GABAergic processing.

fMRS can be used to measure excitatory and inhibitory processing separately within a VOI while participants perform a task (Apšvalka et al., 2015; Ip & Bridge, 2022; Koolschijn et al., 2023; Lally et al., 2014; Mullins, 2018; Stanley & Raz, 2018). However, a major limitation of fMRS is the large VOI size. In this study the VOI consisted primarily of early visual area V1, but the measured glutamate and GABA signals correspond to a mixture between the representations of the RSVP and the annulus with the task-irrelevant coherent motion direction within V1. However, it is likely that the difference in glutamate levels between suprathreshold and near threshold exposure most likely originated from parts of V1 representing the task-irrelevant coherent motion direction rather than the RSVP for the following reasons. First, there was no significant difference in RSVP performance between suprathreshold and near threshold exposure. Second, no significant association was found between initial glutamate levels and subsequent RSVP learning speed. Third, the VOI was placed slightly away from the occipital pole to minimize the inclusion of the foveal confluence with the RSVP representation.

In conclusion, the results of this study suggest that excitatory processing in early visual areas reflecting task relevance and perceptual salience modulates VPL for repeatedly exposed task-irrelevant visual signals. Less pronounced excitatory processing at the earliest stage of cortical visual processing may be an efficient mechanism for reducing VPL for sufficiently salient task-irrelevant visual signals.

## Acknowledgment

We wish to thank Jennifer Lubich and Susanna Hirschle for help with data collection.

---

**Table 1.** MRSinMRS checklist.

|  |  |
| --- | --- |
| **Site (Name or Number):** Brain Imaging Center at University of Regensburg / Bezirksklinikum Regensburg |  |
| **1. Hardware** |  |
| a. Field strength [T] | 3 T |
| b. Manufacturer | Siemens |
| c. Model (software version if available) | Prisma |
| d. RF coils: nuclei (transmit/receive), number of channels, type, body part | 64 channel head/neck coil |
| e. Additional hardware | N/A |
| **2. Acquisition** |  |
| a. Pulse sequence | Siemens MEGAPRESS sequence |
| b. Volume of Interest (VOI) locations | Occipital lobes |
| c. Nominal VOI size [cm<sup>3</sup>] | 2 x 2 x 2 cm |
| d. Repetition Time (TR), Echo Time (TE) [ms] | MEGAPRESS (Edit On & Edit Off): TR 1500 ms; TE = 68 ms |
| e. Total number of Excitations or acquisitions per spectrum | 128 scans in total in each fMRS run |
| f. Additional sequence parameters (spectral width in Hz, number of spectral points, frequency offsets) | Spectral bandwidth: 1200 Hz; Spectral points: 1024 |
| g. Water Suppression Method | WET |
| h. Shimming Method, reference peak, and thresholds for “acceptance of shim” chosen | Automated B<sub>0</sub> field mapping to < 20 Hz |
| i. Triggering or motion correction method | N/A |
| **3. Data analysis methods and outputs** |  |
| a. Analysis software | LC Model Version 6.3-1N |
| b. Processing steps deviating from quoted reference or product | N/A |
| c. Output measure | Tissue-corrected concentrations relative to the composite measure of N-Acetylaspartate and N-Acetylaspartylglutamate |
| d. Quantification references and assumptions, fitting model assumptions | Imported metabolite basis sets included the following spectra. Edit Off: γ-Aminobutyric Acid (GABA), Glutamine (Gln), Glutamate (Glu), Glutathione (GSH), N-Acetylaspartate (NAA), Creatine (Cr), myo-Inositol (mI), L-Lactate (Lac), Aspartate (Asp), Taurine (Tau), N-Acetylaspartylglutamate (NAAG), scyllo-Inositol (Scyllo), Glycerophosphocholine (GPC). Difference: γ-Aminobutyric Acid (GABA), Glutamine (Gln), Glutamate |
|  | (Glu), Glutathione (GSH), N-Acetylaspartate (NAA), N-Acetylaspartylglutamate (NAAG). |
| **4. Data Quality** |  |
| a. Reported variables | CRLB, shimming, S/N |
| b. Data exclusion criteria | CRLB > 25%, shimming > 20 Hz, S/N < 5 |
| c. Quality measures of postprocessing Model fitting | CRLB, S/N |
| d. Sample Spectrum | See Supplementary Figures 5 and 6 |

## Supplementary Materials

**Supplementary Figure 1.**
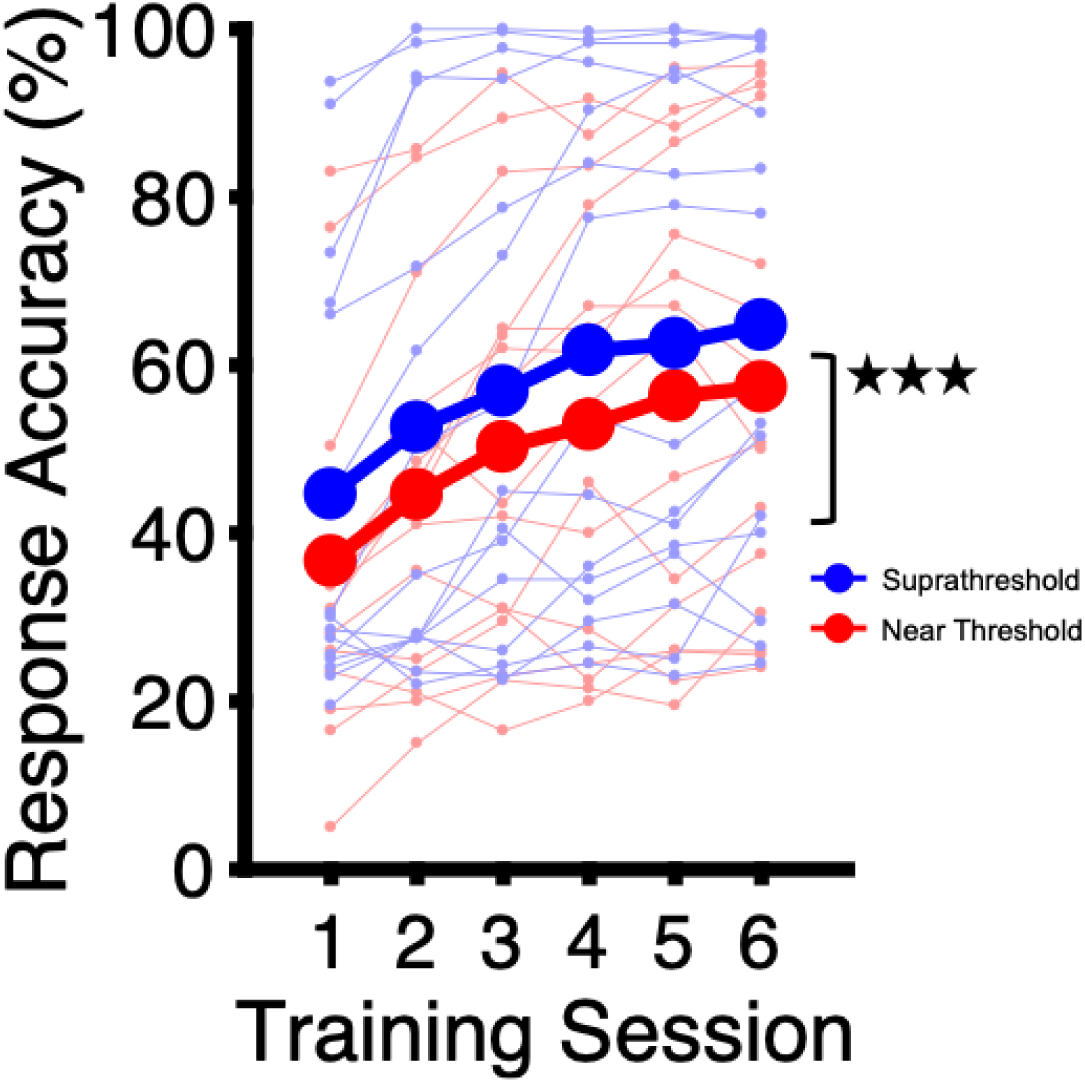
Results for target detection in the RSVP stream in the behavioral experiment. Thick lines show mean response accuracies across participants in each exposure group. Thin lines show response accuracies for each participant. The asterisks show a significant main effect of training session. *** *p* < 0.001.

**Supplementary Figure 2.**
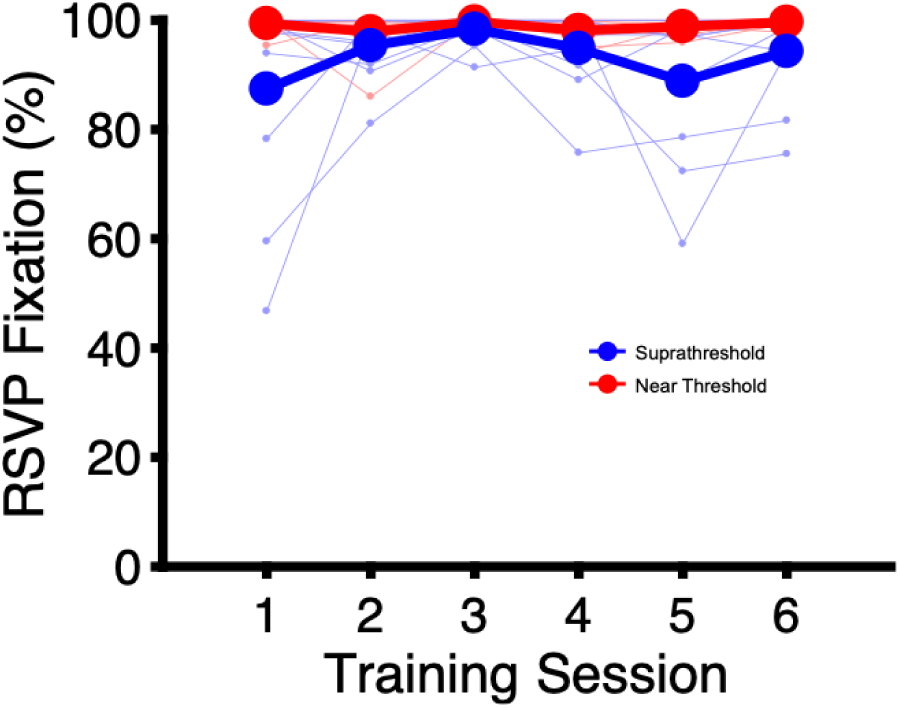
Eye-tracking results in the behavioral experiment. Thick lines show mean percentage of trial time with fixation on the RSVP across participants with eye-tracking results in each exposure group. Thin lines show results for each participant.

**Supplementary Figure 3.**
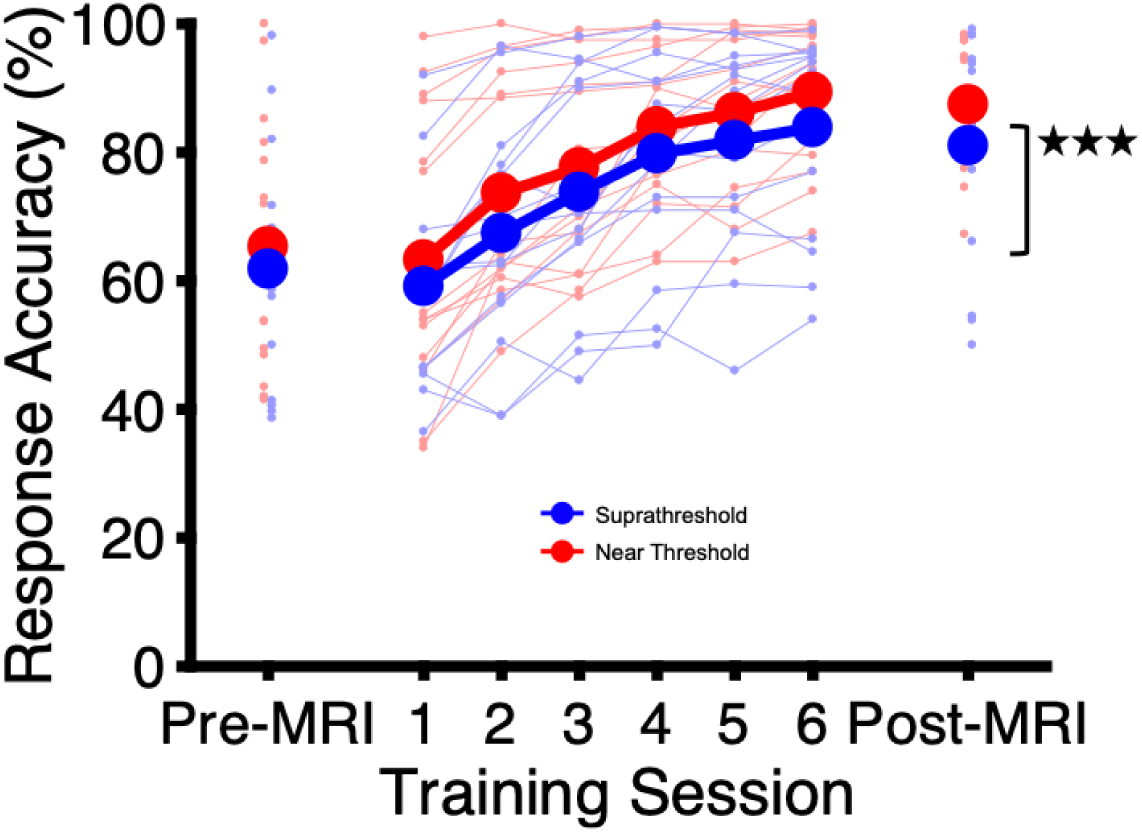
Results for target detection in the RSVP stream in the imaging experiment. The asterisks show a significant main effect of MRI. Otherwise same as Supplementary Figure 1. *** *p* < 0.001.

**Supplementary Figure 4.**
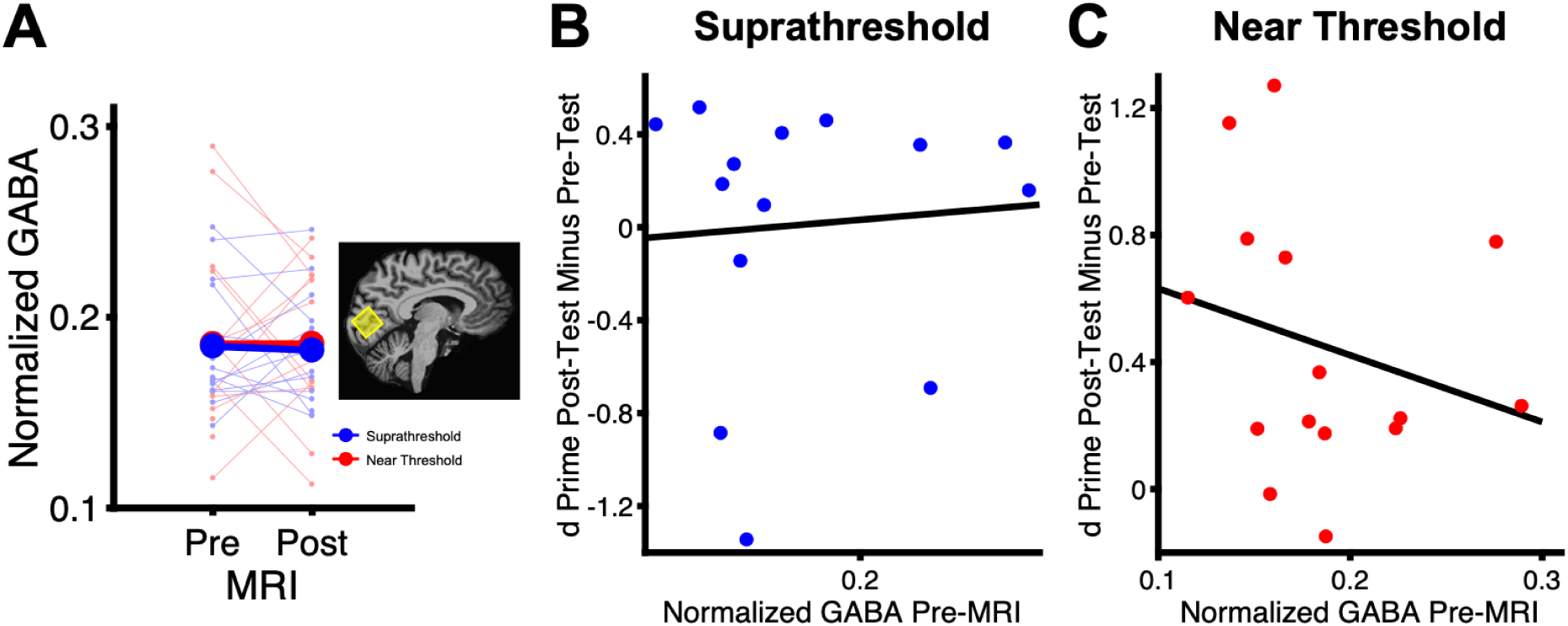
Inhibitory processing levels in early visual areas in the imaging experiment. (**A**) GABA concentration normalized to a control metabolite. Otherwise same as Figure 4A. (**B**) Correlation between GABA concentration in Pre-MRI and sensitivity change for the task-irrelevant learning direction from Pre-Test to Post-Test (see Figure 3B) in the suprathreshold exposure group. Each dot shows the result from a different participant in the suprathreshold exposure group. (**C**) Correlation results in the near threshold exposure group. Otherwise same as (B).

**Supplementary Figure 5.**
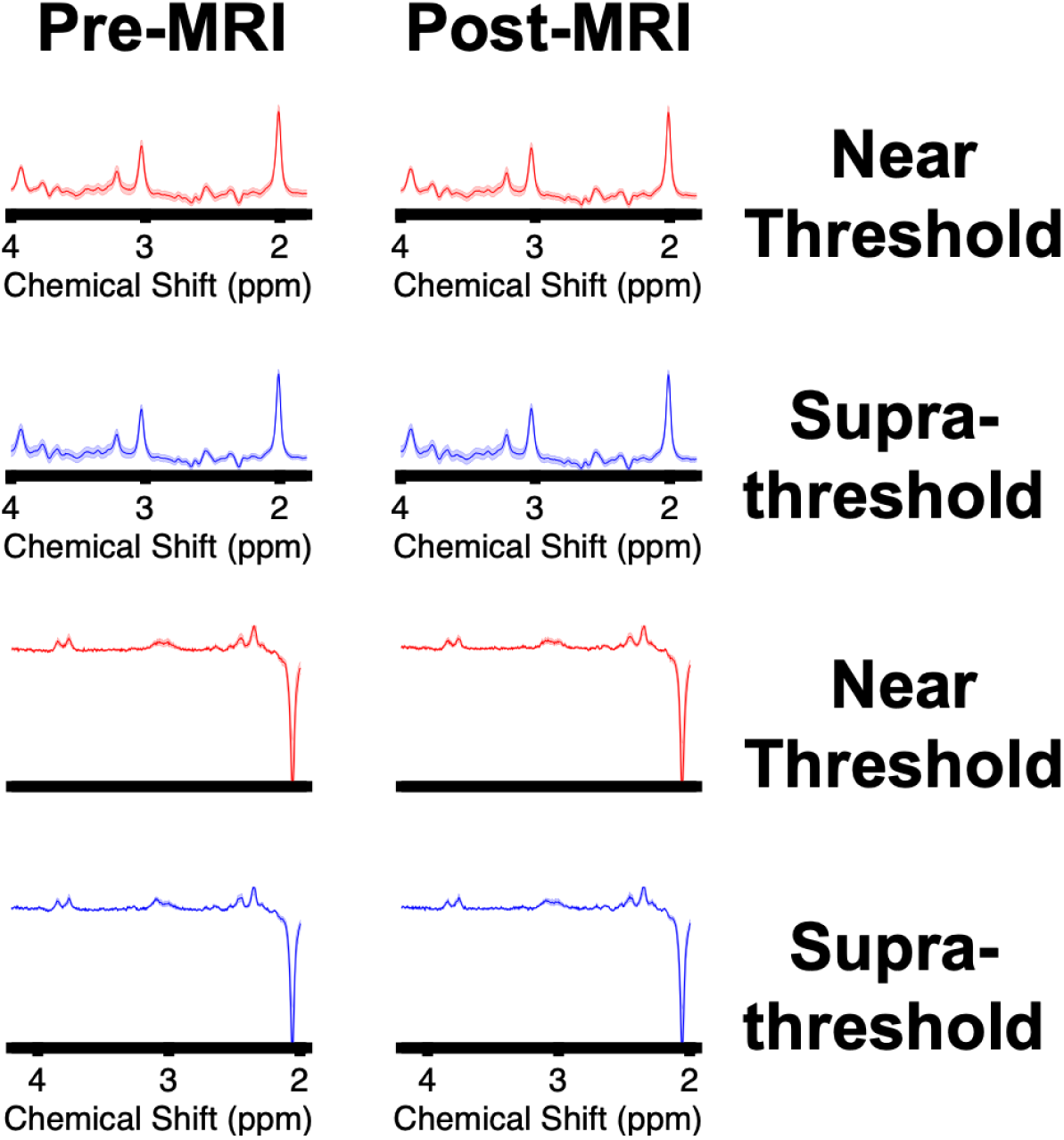
fMRS spectra in the imaging experiment. Top two rows: Edit Off spectrum. The thick line shows the mean spectrum across participants in each exposure group. The shaded area shows standard-error-of-the-mean. Glx between 2.1 and 2.5 ppm and at 3.75 ppm. NAA at 2 ppm. ppm = parts per million of the proton frequency. Bottom two rows: Difference spectrum. GABA at 3 ppm and NAA at 2 ppm.

**Supplementary Figure 6.**
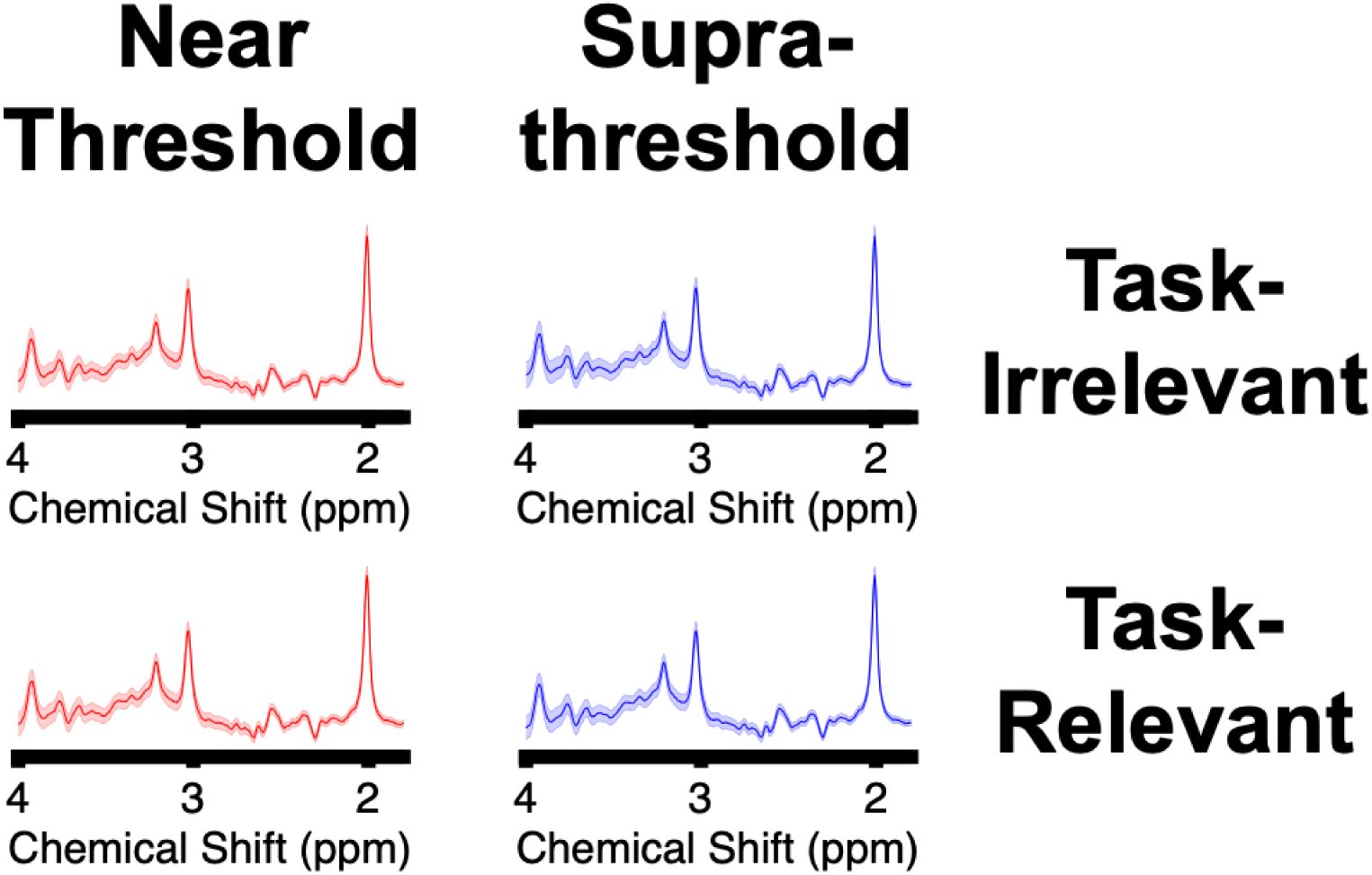
Edit Off spectra in the control imaging experiment. The thick line shows the mean spectrum across participants. The shaded area shows standard-error-of-the-mean. Glx between 2.1 and 2.5 ppm and at 3.75 ppm. NAA at 2 ppm.

